# Calculating and applying pathogen mutational spectra using MutTui

**DOI:** 10.1101/2023.06.15.545111

**Authors:** Christopher Ruis, Gerry Tonkin-Hill, R. Andres Floto, Julian Parkhill

## Abstract

Mutational spectra describe the pattern of mutations acquired during evolution and are driven by factors including mutagens, repair processes and selection. Calculating mutational spectra of pathogen genomic datasets may enable analysis of factors that influence these mutational processes, including replication niches, transmission routes and pathogen biology. Here, we introduce MutTui, which can leverage multiple types of sequence data to calculate and compare mutational spectra of DNA and RNA pathogens. MutTui is available at https://github.com/chrisruis/MutTui.

## Background

Pathogens accumulate mutations during their evolution. These mutations can initially be introduced by polymerase errors [1], natural damage to nucleotides (for example spontaneous deamination of cytosine to uracil [2]), enzyme activity [3,4] or exposure to endogenous or exogenous mutagens [5,6]. Many pathogens encode enzymes and pathways that recognise and repair nucleotide damage [7,8], thereby correcting mutations. Once a mutation has occurred, it is acted upon by selection and genetic drift, which influence whether the mutation persists or is lost. Previous work within the cancer field investigating the mutational patterns associated with these processes has shown that individual mutagens and repair processes drive characteristic mutational signatures [9–15]. These signatures are typically characterised through single base substitution (SBS) spectra that consist of each mutation type (for example C mutating to A, referred to here as C>A) and the nucleotides immediately surrounding the mutation site (the mutational context) [11,12]. Some mutational processes are also associated with double base substitution (DBS) signatures as they induce mutations at adjacent genomic positions [9,13]. Mutational signatures combine to form a mutational spectrum containing the full set of mutations acquired during evolution. Calculation of the mutational spectra of tumours and extraction of their active signatures has enabled inference of the drivers of tumourigenesis [10,11,14–16].

There is growing interest in applying similar analyses to identify active signatures and their drivers within pathogens [17–23]. This may enable inference of endogenous and exogenous factors that influence mutational patterns. For example, pathogens replicate within different niches and exhibit diverse transmission routes, which can result in exposure to different sets of exogenous mutagens. Characterisation and comparison of mutational patterns across pathogens in different niches may identify niche-specific signatures that can subsequently be used to infer niche and transmission pathways for pathogens where this is currently poorly understood. For example, we identified a signature of C:G>A:T mutations associated with the lung across DNA and RNA pathogens that can be used to infer the predominant replication niche(s) of emerging clades of *Mycobacteria* [18] and SARS-CoV-2 [19]. Additionally, gene gains and losses and/or mutations may alter exposure to endogenous mutagens or the activity of repair pathways [23,24]. Comparison of pathogens with different gene contents can therefore identify endogenous signatures that can be used to uncover active cellular pathways in other clades.

Inferring pathogen mutational spectra requires a different approach to calculation of the spectra to that used in cancerous tissues. While it is possible to identify the direction and surrounding context of mutations in a cancerous tissue through comparison with a healthy tissue from the same patient [11,12], such a “healthy tissue” doesn’t exist for pathogen datasets. We have therefore developed MutTui, a novel pipeline to calculate mutational spectra for pathogen datasets. MutTui identifies directional mutations (i.e. inferring the ancestral and derived nucleotides) through ancestral reconstruction onto a pathogen phylogenetic tree. The genetic diversity and rapid evolution of many pathogens means that a single reference genome cannot be used to accurately identify surrounding nucleotide contexts. MutTui solves this problem by updating the reference genome at each node of the phylogenetic tree, thereby identifying the context of each mutation at the time it occurred. Alternatively, MutTui can calculate the mutational spectrum of individual pathogen samples using deep sequencing data and a closely related reference genome. We introduce the application of MutTui to calculate pathogen mutational spectra from multiple sources of input data and present extensions including examination of strand bias and mutational spectra containing only synonymous mutations. We demonstrate the utility of MutTui to examine signatures of DNA repair genes by analysing bacterial hypermutator lineages and show the ability of MutTui to identify mutational signatures associated with replication niches.

## Results

### Overview

MutTui calculates mutational spectra using either an alignment of genetic sequences with an associated phylogenetic tree, or a set of variants from deep sequencing data (**Figure 1**). The methods differ for these two approaches; we first describe spectrum calculation from phylogenetic trees. Here, MutTui initially uses treetime [25] to reconstruct directional mutations (i.e. inferring the ancestral base and the mutated base) onto the phylogenetic tree. These mutations are divided into single mutations (which are not immediately adjacent to another mutation acquired on the same phylogenetic branch), double mutations (pairs of mutations at adjacent genomic positions acquired on the same phylogenetic branch) and tract mutations (three or more mutations at adjacent genomic positions acquired on the same phylogenetic branch). Single mutations are included in the single base substitution (SBS) spectrum; the surrounding context (**Figure S1**) of each mutation is identified using a reference genome that is updated at each node in the phylogenetic tree to incorporate mutations acquired between the root of the phylogeny and that node (**Methods**). The context is therefore inferred within the genome sequence at the start of the respective branch. The direction of mutations on the phylogenetic branches that descend immediately from the root cannot be robustly inferred; by default these branches are excluded by MutTui. Double mutations are incorporated into the double base substitution (DBS) spectrum. Tract mutations are excluded from further analysis. MutTui additionally calculates a mutation type spectrum from the total count of each mutation type summed across contexts.

**Figure 1.**
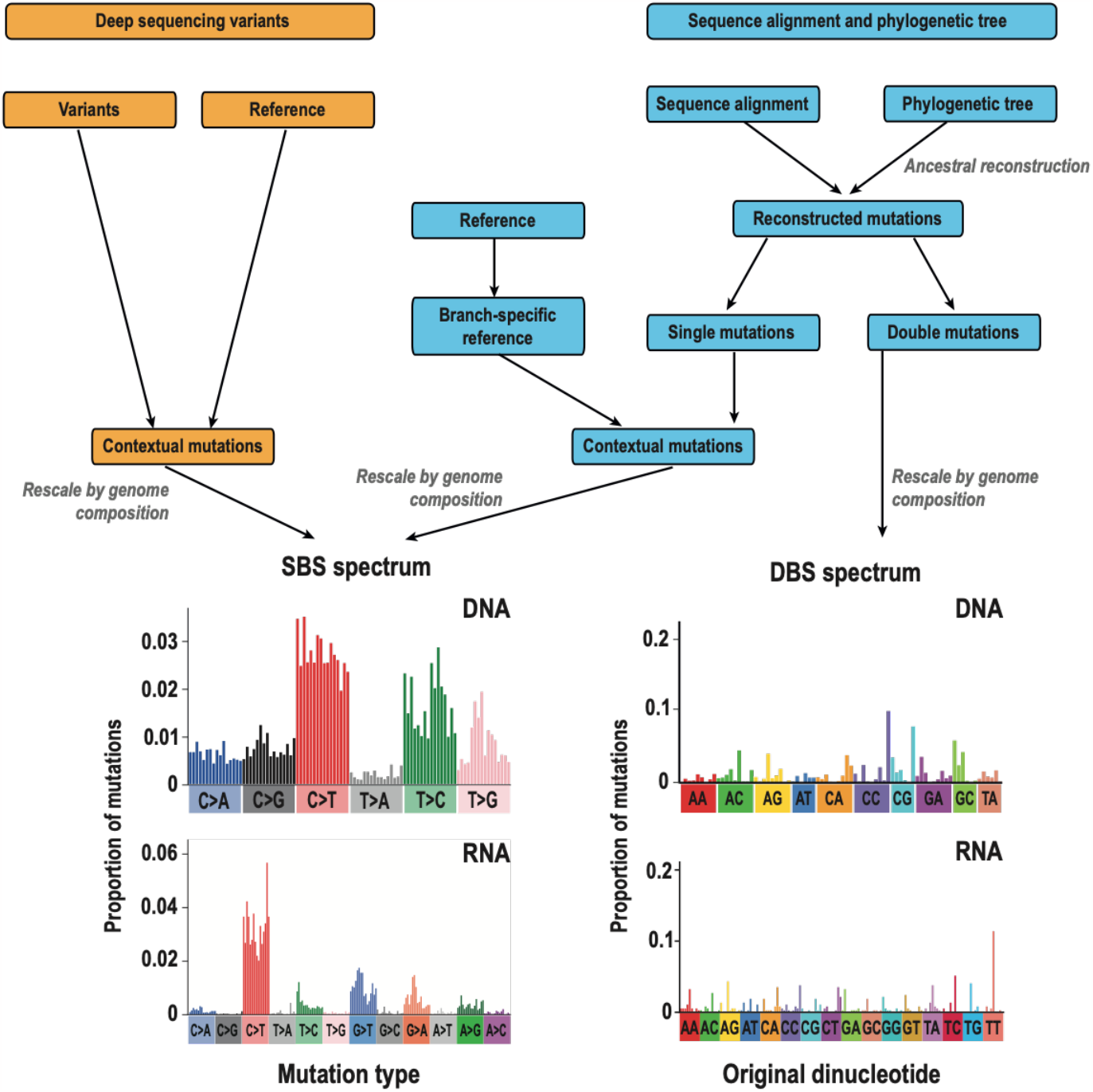
Overview of MutTui pipeline. Spectrum reconstruction can begin with either a sequence alignment and associated phylogenetic tree or a set of variants from deep sequencing data. When using an alignment and tree, mutations are initially reconstructed onto the tree. Mutations are divided into single mutations (not flanked by another mutation acquired on the same phylogenetic branch) and double mutations (pairs of adjacent genomic positions that mutate on the same phylogenetic branch). Tracts of three or more adjacent mutations acquired on the same branch are discarded. The context of single mutations is inferred from a branch-specific reference genome and these contextual mutations are included in the SBS spectrum. Double mutations are included in the DBS spectrum. When using variant data, the context of each mutation is inferred from a closely related reference genome (typically the genome that was mapped against). In all cases, mutations are rescaled by genome composition to produce SBS and DBS spectra that are comparable across datasets. The pipeline proceeds with the same steps for DNA and RNA datasets but with different outputs; in DNA datasets, reverse mutations cannot be distinguished so are combined resulting in 96 SBS mutations (16 contexts within each of six mutation types) and 78 DBS mutations. In RNA datasets, reverse mutations are not combined resulting in 192 SBS mutations (16 contexts in each of 12 mutation types) and 144 DBS mutations.

Recombination may result in a subset of mutations being incorrectly inferred to occur on multiple phylogenetic branches and/or incorrect inference of mutation direction, and can therefore cause aberrant spectra. It is therefore currently essential that recombination is removed prior to spectrum calculation and datasets should therefore be chosen to produce phylogenetic trees that are shallow enough to enable recombination removal while being diverse enough to contain at least several hundred mutations. While many datasets will contain whole genome sequences, the input alignment can also contain only individual genes or genomic regions for rapidly evolving pathogens.

To enable comparison of mutational patterns between pathogens with different genomic compositions (and therefore different opportunities for mutation), SBS and DBS spectra are rescaled by genome composition (**Methods**). This rescales mutation counts to comparable counts per available site. MutTui also outputs a mutation type spectrum containing counts of each mutation type summed across contexts.

To calculate a mutational spectrum from a deep sequencing sample, MutTui uses variant data (which can be calculated through any other programme chosen by the user) and the reference genome that was mapped against (**Figure 1**). The direction of each variant is inferred to be from the reference base to the variant base and the surrounding context of the mutation is inferred from the reference genome. Therefore, the chosen reference should be as closely related to the sample as possible. Alternatively, in cases where a strong transmission bottleneck can be assumed and where most mutations won’t become dominant, the reference genome can be constructed using the dominant nucleotide at each genome position. As variant data typically lacks linkage between variants, each mutation is treated independently and included in the SBS spectrum.

### DNA vs RNA spectra

MutTui can calculate mutational spectra for both DNA and RNA pathogens, which have different replication processes. In DNA genomes, replication of both strands proceeds simultaneously. Mutations that arise during synthesis in one strand will be copied onto the other strand during replication. Opposite mutations occurring in different strands can therefore appear as the same mutation in the sequenced consensus genome (**Figure S2**). For example, a C>T mutation in one strand will be read equivalently to a G>A mutation in the other strand. We therefore combine symmetric mutations in DNA spectra, as is common practice in cancer mutational spectra [11,12]; SBS spectra of DNA pathogens consist of 96 contextual mutations (16 surrounding contexts within each of six mutation types) and DBS spectra contain 78 double substitutions (**Figure 1**) [9,11–13].

However, single stranded RNA virus genomes exist as either a positive or negative RNA strand. The genome is maintained in this strand outside of active replication, which occurs through an RNA intermediate of the opposite polarity. Mutations that occur during replication will be symmetric, as in DNA. However, mutations arising outside of replication will predominantly affect the genomic strand and therefore will not be symmetric. Correspondingly, previous studies have shown significant differences between symmetric mutations in RNA pathogens [19,21] and we therefore maintain mutations on each strand separately in RNA spectra. SBS spectra of RNA pathogens therefore consist of 192 contextual mutations (16 surrounding contexts within each of 12 mutation types) and DBS spectra consist of 144 double substitutions (**Figure 1**).

Pathogens whose genomes spend the majority of the time as double stranded molecules (for example double strand RNA virus) should be treated as DNA pathogens for spectrum calculation while pathogens whose genomes spend the majority of the time as single stranded molecules (for example single stranded DNA viruses) should be treated as RNA pathogens.

### Coding gene information enables calculation of synonymous SBS spectra and examination of coding strand bias

MutTui can optionally be provided with a GFF file containing gene coordinates within the reference genome. This enables calculation of mutational spectra containing only synonymous mutations (**Figure S3**) and, for DNA pathogens, a coding strand-specific SBS spectrum in which mutations are subdivided based on whether the starting pyrimidine base is on the transcribed or untranscribed strand (**Figure S4**). Synonymous mutation-only spectra may be useful to minimise the impact of selection on the observed mutational patterns while coding strand-specific spectra may enable further investigation of mutagens that exhibit coding strand bias due to the action of transcription coupled nucleotide excision repair [9,10,26].

### Branch labelling

Comparing spectra between different sets of phylogenetic branches can identify the mutational patterns and processes that differ between groups. Where the groups to be compared are separate monophyletic clades, their mutational spectra can be calculated through separate MutTui runs. However, it is often useful to compare spectra between sets of branches within a single phylogenetic tree based on some associated metadata, which may change once or many times across the tree. Such comparisons are enabled by MutTui through assigning each phylogenetic branch a label and calculating a mutational spectrum across branches with the same label (**Figure 2**). MutTui can label the phylogenetic tree when given a root state and the branches across the tree on which the label changes. This tree can then be provided to calculate spectra for each label. Alternatively, MutTui can infer locations of label changes using discrete trait reconstruction through treetime [25]. We would recommend user labelling of the tree where possible.

**Figure 2.**
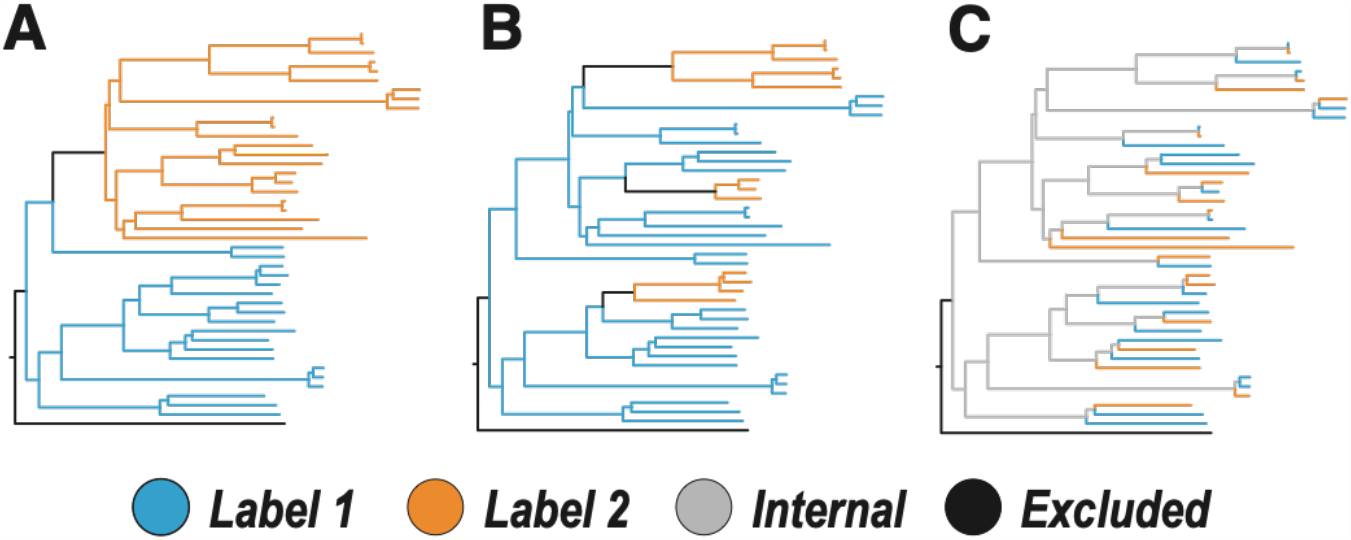
Examples of branch labelling. In each example, we aim to compare the mutational patterns of label 1 (blue) with label 2 (orange). Branch colours indicate the category to which each branch is assigned. Branches that descend immediately from the root are excluded in each case. In **A** and **B**, branches on which the label changes are also excluded. **A** shows a single label change whose location within the phylogenetic tree can be robustly inferred. In **B**, there are multiple label changes but the location and direction of label changes can be robustly inferred. (**C**) The label changes regularly such that the label of internal branches cannot be robustly inferred. In this case, we can examine mutational patterns on the tip phylogenetic branches only and the internal branches are assigned to a separate “internal” category. To enable comparison in this case, the branches on which the label changes are included in spectrum calculation.

In most cases, the branches on which a metadata label changes can be inferred but the exact time of the change along the branch cannot. Mutations therefore cannot typically be accurately assigned to a label on branches where the label changes; MutTui excludes mutations on such branches by default. Where patterns of metadata are complex and/or internal phylogenetic branches cannot be assigned to a label with certainty, we recommend labelling the tip phylogenetic branches but excluding internal branches by assigning them to an ‘internal’ label (**Figure 2**).

### Comparing mutational spectra

MutTui includes several methods to compare mutational spectra. The overall similarity between a set of mutational spectra can be quantified by calculating the cosine similarity [11] between all pairs. The relationships between groups of mutational spectra can be analysed by principal component analysis. Depending on the question of interest, this clustering can be based on overall spectrum composition, mutation type proportions, context proportions within an individual mutation type or the cosine distances between spectra.

Enriched mutations and/or mutation types can be identified through spectrum subtraction where the proportion of each mutation in one spectrum is subtracted from that in the other spectrum. The significance of differences between spectra can be assessed through a permutation test where the difference in mutation proportions in the real data is compared with the distribution of differences across 1000 random permutations of the mutations across spectra [23].

MutTui can also compare contextual mutation patterns within individual mutation types across different spectra (**Figure S5**). Here, contextual mutation proportions are regressed and the Pearson’s correlation coefficient of this regression is compared with the distribution of correlation coefficients across 1000 randomisations of the contextual proportions within each spectrum.

Confidence intervals on the proportion of individual mutations and mutation types can be calculated by assuming a beta-binomial distribution and employing the Wilson score interval using the total number of mutations as the number of trials and the mutation proportion as the success proportion.

### Sensitivity analysis

To examine the reliability of the mutational spectra calculated by MutTui, we simulated 100 datasets each containing approximately 10,000 mutations sampled from the mutational spectrum of *Mycobacterium tuberculosis* lineage 4 we had calculated previously [18] (**Figure S6**). For each simulation, we compared the mutational spectrum calculated by MutTui with the real sampled mutation counts. In 91 of the 100 simulations, the real and calculated mutational spectra were identical. In the remaining nine simulations, MutTui inferred one extra mutation out of ∼10,000, likely due to the ancestral reconstruction inferring a single mutation to have occurred on two separate branches.

Robust estimation of the mutational spectrum requires the dataset to contain sufficient mutations. We assessed the required number of mutations by downsampling multiple DNA and RNA SBS spectra and calculating the cosine similarity between the original and downsampled spectra (**Figure 3**). Across these datasets, an accurate representation of the SBS spectrum (cosine similarity > 0.95) can be achieved with at least 600 mutations (range 300-600 mutations depending on the dataset, **Figure 3**).

**Figure 3.**
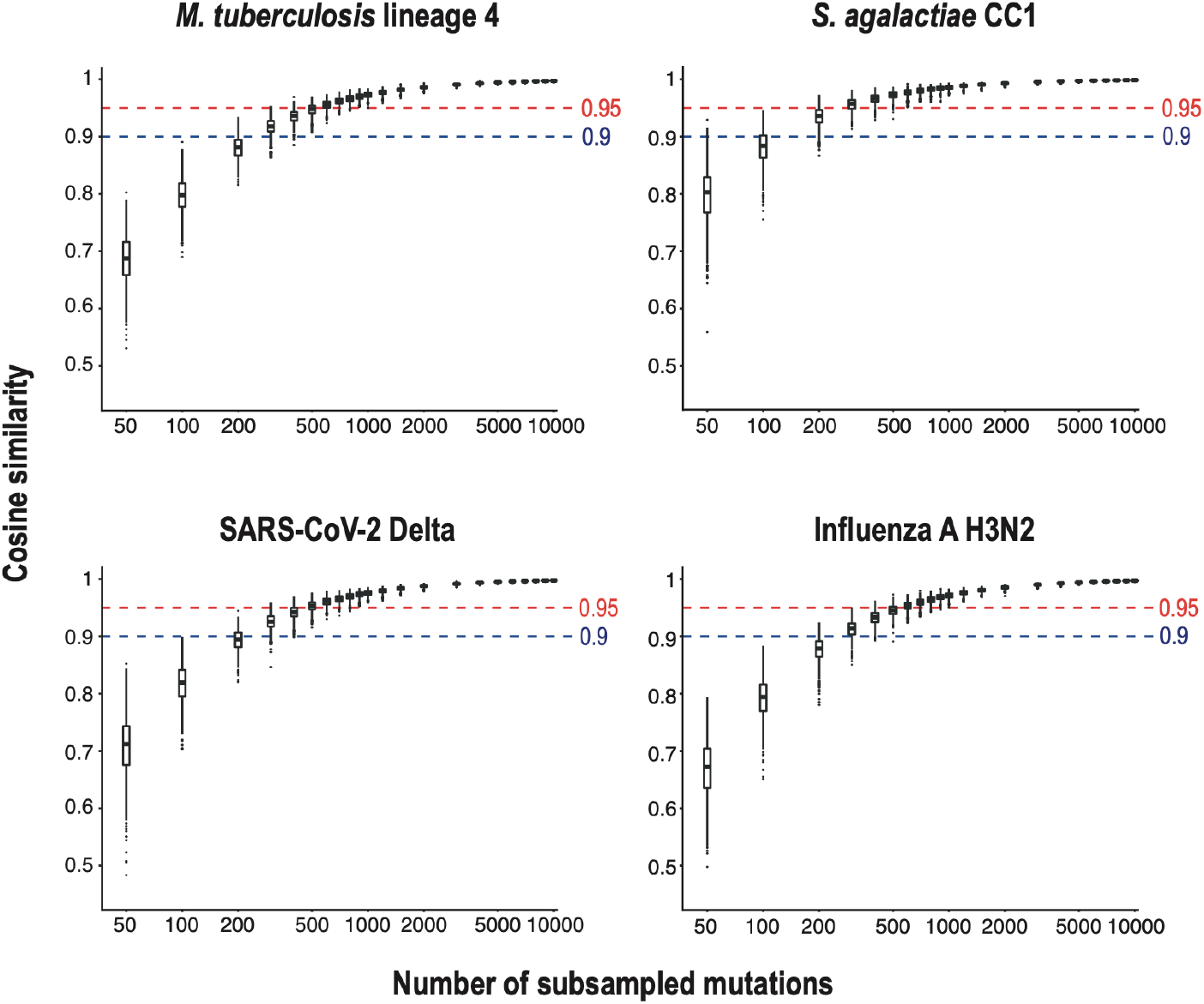
Assessment of the minimum number of mutations required to accurately calculate the SBS spectrum. We converted two DNA SBS spectra (*M. tuberculosis* lineage 4 and *S. agalactiae* CC1) and two RNA SBS spectra (SARS-CoV-2 Delta and Influenza A H3N2) to contain 10,000 mutations. These spectra were downsampled to different numbers of mutations and the cosine similarity to the original spectrum was calculated. The number of mutations required to accurately represent the original spectrum was the minimum with median cosine similarity > 0.95.

To test the robustness of the calculated SBS spectrum to tree topology, we ran MutTui on ten different phylogenetic trees sampled from a posterior distribution for a DNA dataset and an RNA dataset [18,19]. The inferred SBS spectrum was highly similar with all topologies (cosine similarity >0.99 between each spectrum pair, **Figure S7**). Additionally, the confidence intervals on the proportion of each SBS mutation and each mutation type overlapped in each run. Calculated mutational spectra are also highly similar (cosine similarity >0.99) when using either the GTR or HKY substitution model for ancestral reconstruction on the same maximum likelihood phylogenetic tree.

The choice of root location within the phylogenetic tree can alter the direction of a subset of mutations and therefore might alter the calculated spectrum. We recommend using the most robust rooting strategy available for a dataset, which might use an outgroup if a closely related sequence can be identified or employ temporal information through a time tree, or use a root that maximises root-to-tip correlation [27]. SBS spectra calculated with different phylogenetic rooting strategies (outgroup rooting, rooting to maximise root-to-tip correlation and midpoint rooting) were highly similar (cosine similarity >0.99, **Figure S8**), suggesting that any of these strategies may enable robust spectrum calculation.

Most of the potential incorrect inferences caused by the position of the root will likely fall on the two branches descending immediately from the root. MutTui therefore does not use these in reconstructing mutational spectra.

### Identifying mutational signatures associated with DNA repair genes from natural hypermutator lineages

To demonstrate the utility of MutTui to examine signatures of DNA repair processes, we use extraction of *nth* gene signatures as an example. The product of the *nth* gene (also called Endonuclease III) repairs multiple types of damaged nucleotides, including oxidised pyrimidines and uracil incorporated in DNA [28]. We previously calculated *nth* signatures using different data sources from two species: hypermutator lineages within a phylogenetic tree of *Mycobacterium leprae* and variant data from a chronic *Mycobacterium abscessus* infection [18]. The *M. leprae* phylogenetic tree consists of five independent lineages leading to six sampled sequences with evidence of *nth* mutation causing hypermutation [29]. To calculate the mutational spectrum of these hypermutators, we used MutTui to label phylogenetic branches into two categories: branches that are not downstream of *nth* mutation were labelled B (for background) while branches with evidence of *nth* hypermutation were labelled *nth* (**Figure 4A**). We then ran MutTui using this labelled tree to calculate a SBS spectrum of the mutations on the *nth* branches (**Figure 4B**).

**Figure 4.**
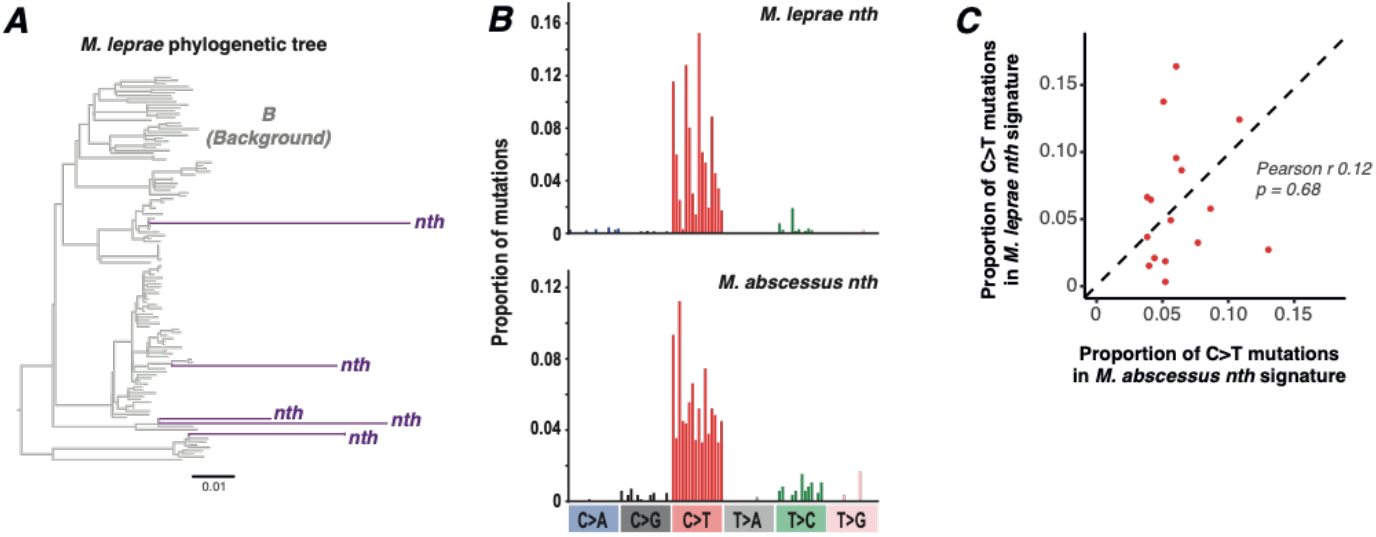
Extraction of *nth* gene signatures. (**A**) Phylogenetic tree of *M. leprae* with branches coloured by their label for MutTui. Grey branches do not exhibit *nth* hypermutation so are labelled B (for background). Five separate branches have acquired *nth* mutations and exhibit hypermutation; these branches are coloured purple. The *M. leprae nth* signature was identified by combining mutations acquired on the branches coloured purple. (**B**) Mutational spectra are shown for *nth* hypermutator lineages in *M. leprae* (calculated from the branches coloured purple in panel **A**) and *M. abscessus* (calculated from SNPs acquired during chronic infection after mutation of *nth*). Each signature predominantly exhibits C>T mutations. (**C**) Comparison of contextual proportions within C>T mutations between the *M. leprae* and *M. abscessus nth* signatures. P-value calculated through comparison of the Pearson’s r correlation in the real data with the distribution across 1000 randomisations of contextual proportions within each dataset.

To calculate the *M. abscessus nth* signature, we obtained previously published variant data from a *nth* hypermutator lineage isolated from a chronic lung infection [30]. We calculated the SBS spectrum of these variants using MutTui, employing the closely related reference (that the reads were mapped against) to infer surrounding contexts (**Figure 4B**).

The resulting SBS spectra are both dominated by C>T mutations (**Figure 4B**). These represent the mutations that are typically repaired by the product of *nth*. Comparing the signatures with MutTui shows different contextual patterns within the C>T mutations (**Figure 4C**), suggesting either different contextual patterns of DNA damage and/or differential context-specific repair by other proteins.

### Extraction of mutational signatures associated with lung and environmental niches

We use the extraction of lung and environmental mutational signatures as an example of applying MutTui to examine niche-specific signatures [18]. In this example, there are multiple independent transitions from predominantly environmental clades causing opportunistic infections to clades specialising in infecting and transmitting through the lung niche within *Mycobacteria* and *Burkholderia*. By calculating and comparing mutational spectra across these clades, we can identify any signatures that are acquired upon independent niche switches. In most cases, we calculated SBS spectra for individual clades that can be assigned to a single niche, for example *Mycobacterium avium* can be assigned to the environment and *Mycobacterium tuberculosis* lineage 4 to the lung (**Figure 5A**). For these clades, we ran MutTui without branch labelling to calculate the SBS spectrum of the complete clade. However, in several cases, isolates with the ability to replicate in different niches are present in the same phylogenetic tree and we therefore needed to label branches to calculate niche spectra. We calculated the *Mycobacterium canettii* spectrum using a previously published phylogenetic tree [30] that also contains *M. tuberculosis* samples. Here, we labelled the tree into “*M. canettii*” and “*M. tuberculosis*” regions and calculated the spectrum of the “*M. canettii*” branches (**Figure 5A**). A similar labelling strategy was used for *Mycobacterium kansasii* (**Figure S9**), as the phylogenetic tree can be divided into the *M. kansasii* main cluster (MKMC) and non-main cluster [31]. We calculated the spectrum of the non-main cluster which is environmental, but excluded the MKMC branches as the niche of this clade is poorly understood [18]. Finally, we calculated SBS spectra for environmental *Mycobacterium abscessus* clades. Here, we excluded the dominant circulating clones (which likely predominantly replicate within the human lung [18,32]) and, due to the high level of diversity within the species, divided the remaining samples into FastBAPS clusters [33] (**Figure S10**). We calculated an SBS spectrum for each FastBAPS cluster separately and then combined the calculated spectra [17].

**Figure 5.**
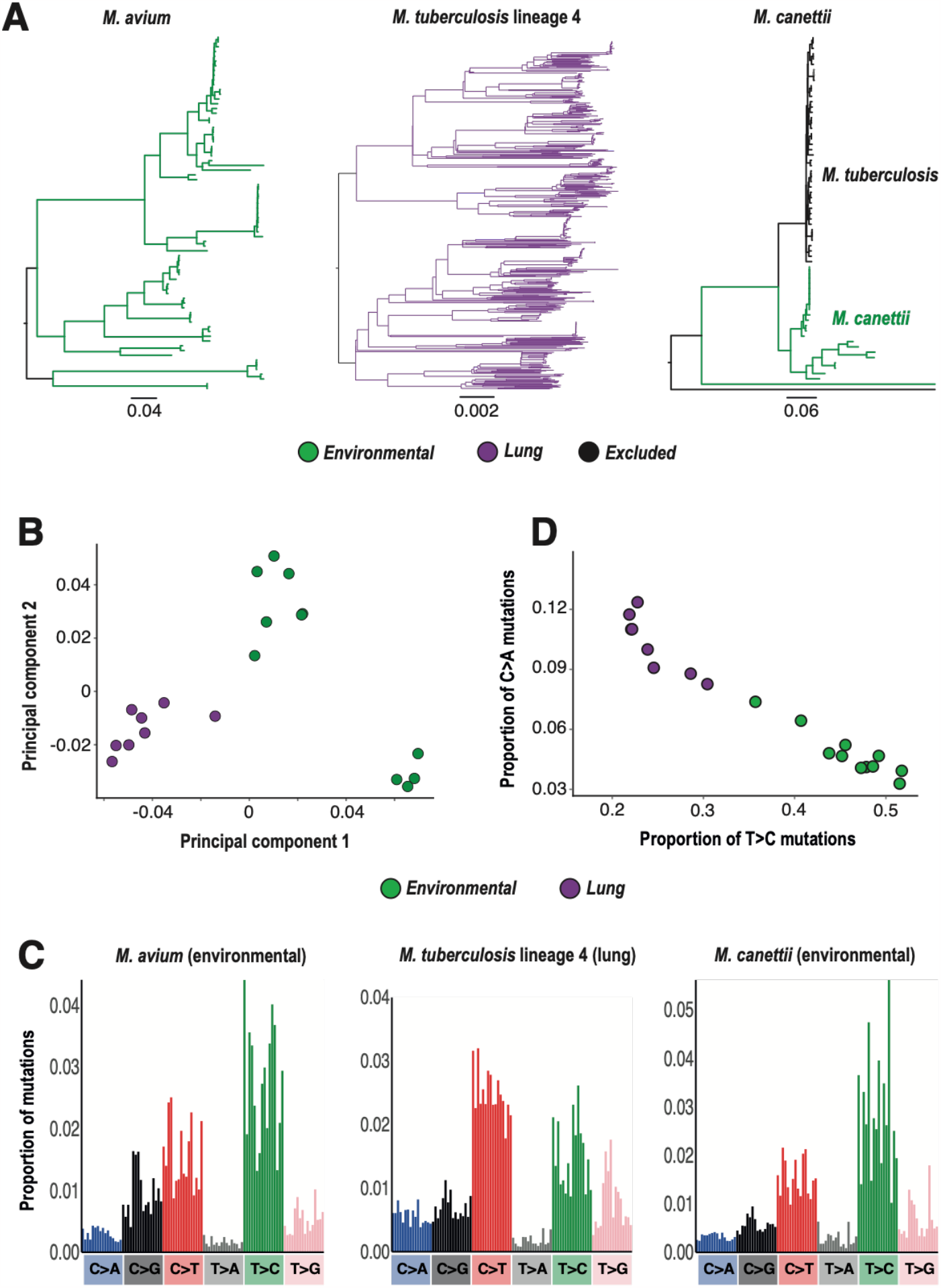
Comparison of SBS spectra from lung and environmental *Mycobacteria* and *Burkholderia*. (**A**) Examples of *Mycobacteria* phylogenetic trees with branches labelled by niche. *M. avium* is an environmental organism while *M. tuberculosis* lineage 4 is a lung pathogen so a single label is applied to all of the branches within these clades. The *M. canettii* spectrum was calculated from a phylogenetic tree that also includes *M. tuberculosis* isolates. Therefore to calculate the *M. canettii* spectrum, we examined branches prior to the divergence of *M. tuberculosis*. In each tree, branches that descend directly from the root node are excluded. Scale bars show expected number of substitutions per variable site. (**B**) Principal component analysis based on composition of SBS spectra coloured by niche. (**C**) SBS spectra for the clades shown in panel **A**. Each spectrum is rescaled by genomic composition. (**D**) Comparison of the proportion of C>A and T>C mutations across SBS spectra coloured by niche. T>C mutations are consistently elevated in environmental clades while C>A mutations are consistently elevated in lung clades.

For the signature discovery, we only included clades whose niche is well characterised, and therefore excluded the *M. abscessus* DCCs and *M. kansasii* MKMC. Principal component analysis of the calculated SBS spectra showed a separation between lung and environmental clades (**Figure 5B**), indicating that one or more mutations differs between niches. Visual inspection of the calculated SBS spectra suggested higher C>A mutations in the lung and higher T>C mutations in the environment (**Figure 5C**). Plotting mutation proportions confirmed this across SBS spectra (**Figure 5D**), indicating that the level of C>A and T>C mutations may be used to infer the predominant niches of other *Mycobacteria* and *Burkholderia* clades.

## Discussion

Mutational spectra can provide novel and important insights into pathogen niches, transmission routes and biology. We have developed MutTui to enable calculation of robust mutational spectra for pathogen datasets, using either a sequence alignment and associated phylogenetic tree or variants from deep sequencing data. MutTui includes several methods to compare mutational spectra between closely related or diverse pathogens calculated across multiple trees or through branch labelling within a single tree. We have demonstrated how MutTui can be employed to calculate DNA repair signatures and extract mutational signatures associated with different niches. While we employed these as illustrative examples, MutTui can be used to investigate any factors that influence mutational patterns. Furthermore, MutTui can provide valuable insights when combined with real time pathogen surveillance, for example to identify movements of a pathogen into a new niche which may be associated with alternative virulence or transmission dynamics [19].

MutTui contains several post-processing scripts to facilitate downstream analyses, including splitting a calculated spectrum into spectra for different groups of branches and calculating summary statistics for each phylogenetic branch.

It is currently essential to remove recombination prior to spectrum calculation. For rapidly evolving RNA viruses with high recombination rates, it may be possible to run MutTui on single genes or genomic regions between recombination hotspots. In future, we plan to investigate if it may be possible to examine datasets containing recombination by collapsing reverse mutations within the spectrum.

MutTui is written in python (versions 3.8+) and is available under the open source MIT license at https://github.com/chrisruis/MutTui. The raw alignments, phylogenetic trees, reference genomes, deep sequencing variants and scripts used in the analyses described above are available at https://github.com/chrisruis/MutTui_manuscript. In the future, we plan to extend MutTui, including through the use of substitution rates rather than proportions in the spectra, development of tools to extract and assign mutational signatures within pathogen datasets and incorporate further post-processing tools.

In summary, we have developed a set of software tools to extract mutagenic signatures from bacterial and viral datasets. We have given examples where this has enabled the identification of replication niches, and the effects of repair systems. We envisage a growing set of investigations that such analyses can be used for in the future.

## Methods

### MutTui algorithm

MutTui can calculate pathogen spectra using two types of input data: a sequence alignment with an associated phylogenetic tree, or variants from deep sequencing data. In either case, MutTui infers directional mutations and, to produce SBS spectra, identifies the surrounding nucleotide context of each mutation. When working with RNA datasets, thymine is used in place of uracil throughout, as is common practice with RNA sequences.

When provided with a sequence alignment and phylogenetic tree, “GTR --infer” is used as default during ancestral reconstruction with treetime and will infer the substitution model that best fits the data. A different substitution model can be specified, as can any other treetime parameters.

To infer the surrounding context of each mutation within the SBS spectrum, MutTui uses a reference sequence and a file that specifies the conversion from alignment position to position in the reference (this file can be generated with MutTui and is not required if the alignment contains all of the positions that are in the reference). The reference sequence is updated at each branch in the tree to incorporate mutations acquired between the root of the tree and the start of the branch. Here, the reconstructed base at the root of the tree is inserted into the reference at each alignment position to construct the root ancestor. At each downstream branch, the mutations acquired between the root and the start of the branch are inserted into the root ancestor to produce a “branch ancestor” sequence. MutTui infers the context of each mutation as the immediately adjacent nucleotides within this branch ancestor and thereby infers the contexts within the genomic background of the branch on which the mutation occurred. The direction of mutations that occurred on the branches that descend immediately from the root cannot be reliably inferred; these branches are excluded from spectrum calculation by default.

Mutations that occur between positions given ambiguity codes (i.e. not A, C, G or T) and, for the SBS spectrum, mutations that have at least one surrounding position with an ambiguity code, are excluded. Additionally, mutations at the first or last position of the sequence are excluded.

To enable comparison of mutational patterns across pathogens with different nucleotide and triplet compositions, MutTui rescales the calculated spectra to convert from counts to comparable counts per available site. Here, the count is divided by the number of the starting triplet (for SBS spectra) or starting dinucleotide (for DBS spectra) in the reference genome.

To enable easy comparison of the mutational patterns on individual phylogenetic branches, MutTui includes a “branch specific” option that will calculate spectra for each individual branch that contains more than a user-specified number of reconstructed mutations.

MutTui can calculate synonymous and strand bias spectra through the use of a GFF file containing gene coordinates within the provided reference sequence. To calculate a synonymous spectrum, the coding effect of each reconstructed mutation is investigated by inserting the mutations on a branch into the “branch ancestor” sequence. All mutations on the branch are inserted together to enable multiple mutations within the same codon to be examined together. The coding effect of each mutation is identified and synonymous mutations extracted. To calculate the strand bias spectrum for DNA pathogens, each coding mutation is separated based on whether the starting pyrimidine base is on the coding or noncoding strand (for example, in C:G>A:T mutations by whether the cytosine nucleotide is on the coding or noncoding strand) to produce a 192 base SBS spectrum.

To calculate mutational spectra from deep sequencing samples, MutTui does not call variants but can use variants from a VCF file. Alternatively, a flexible format variant file can be provided which can have any format as long as it includes the columns: position (containing the position of the variant in the reference), reference (containing the reference nucleotide at the position) and variant (containing the variant nucleotide at the position). Each provided variant is assumed to occur once; if variants have likely occurred on multiple occasions, they should be provided as multiple entries in the variant file. As linkage between variants is typically absent in variant data, a DBS spectrum is not calculated and all mutations are included in the SBS spectrum.

Mutational spectra calculated from variant data might be useful to examine, for example, mutations acquired during individual chronic infections. As individual samples often won’t contain many variants, it may be necessary to combine multiple samples, either through generating a single combined variant file prior to running MutTui or combining multiple calculated spectra.

### Branch labelling

MutTui calculates a SBS spectrum, a DBS spectrum and a mutation type spectrum for each label included within the phylogenetic tree. By default, all branches are assigned the same label. The tree can alternatively be subdivided into two or more labels through branch labelling, which is a two stage process. In the first stage, the tree is labelled to give all internal nodes a node number. Tip branches retain the name of the tip sequence. This tree can be viewed with a tree viewing program such as FigTree [34] to identify the node numbers and tip names of branches where the label should change. To add the final labels to the tree, MutTui takes a root label and combinations of branch names where the label changes and what the label changes to. MutTui will label the tree so branches inherit the label of their parental branch unless the branch is specified with a label change.

Alternatively, if the branches on which the label changes are unknown and difficult to identify manually, MutTui can infer the location of label changes employing the mugration model within treetime [25]. Here, an additional metadata file is provided with the label of each tip sequence. In rare cases, there may be multiple equally supported ancestral states at the root of the tree; to resolve this, a desired root label can be provided.

### Output

MutTui outputs multiple spectra files for each run. For each phylogenetic branch label, MutTui outputs a SBS spectrum, the SBS spectrum rescaled by genomic composition, a DBS spectrum and a mutation type spectrum. MutTui also outputs “all included mutations” files for the SBS and DBS spectra summarising the included mutations and a “mutations not included” file summarising the mutations that were inferred by treetime but not included in spectrum calculation.

### Post-processing

MutTui comes packaged with several post-processing scripts for spectrum analyses. The comparison methods discussed above are run post-spectrum calculation. A branch-specific summary of mutations can be generated in which statistics including the total number of mutations and the number and proportion of each mutation type is calculated for each branch. The mutations from a single MutTui run can be split into any number of branch labels by providing a file containing branch names and their corresponding labels. This may be useful to compare many different label sets on a single tree.

To enable further investigation of the mutation types that differ between spectra, MutTui can plot comparisons of mutation type proportions and calculate ratios of each mutation type pair between spectra.

### Sensitivity analysis

To test the reliability of the mutational spectra calculated by MutTui, we carried out simulations using the *M. tuberculosis* lineage 4 phylogenetic tree and SBS spectrum we calculated previously [18] (**Figure S6**). We rescaled the branch lengths in the tree to a total branch length of 10,000 and then iterated through the tree; for each branch, we sampled a random number from a Poisson distribution with lambda equal to the branch length and chose this number of mutations randomly from the *M. tuberculosis* lineage 4 SBS spectrum. This results in roughly 10,000 mutations across the tree. Tip sequences are built up of triplets; for each sampled mutation, the tip is assigned the ancestral triplet if it is not downstream of the branch on which the mutation occurred, or the mutated triplet if it is downstream of the branch (**Figure S6**). We record the number of each SBS mutation that is sampled. We ran MutTui on the resulting alignment with the original tree and the calculated SBS spectrum was compared with the mutation counts sampled during the simulation. 100 simulations were run and the respective SBS spectra compared.

To assess the minimum number of mutations required to robustly estimate a spectrum, we used previously calculated SBS spectra of two DNA and two RNA pathogens: *M. tuberculosis* lineage 4, *Streptococcus agalactiae* CC1, SARS-CoV-2 variant Delta and Influenza H3N2 [18,19]. These spectra were scaled to contain 10,000 total mutations. We sampled this 10,000 mutation spectrum to different levels with replacement and compared the sampled spectrum to the original using cosine similarity. 1000 downsamplings were carried out at each of 50, 100, 200, 300, 400, 500, 600, 700, 800, 900, 1000, 2000, 3000, 4000, 5000, 6000, 7000, 8000, 9000 and 10,000 mutations. We considered the sampled spectrum to be a good representation of the original spectrum when median cosine similarity was greater than 0.95.

We assessed sensitivity of the reconstructed spectra to tree topology by running MutTui on ten different tree topologies sampled from a posterior distribution of trees for *S. agalactiae* CC1 (DNA) and MERS coronavirus (RNA). The posterior distribution was calculated by running MrBayes v3.2.7a [35] on our previously published sequence alignments [18,19] using the GTR model of nucleotide substitution with gamma site heterogeneity and four gamma classes. Two independent MCMC chains were run for at least 1,000,000 steps and convergence (defined as all ESS values over 200) assessed using Tracer v1.7.2 [36]. Ten phylogenetic trees were sampled from the posterior distribution; MutTui was run on each phylogenetic tree with the same alignment, position conversion file and reference. The proportion of each SBS mutation was calculated and compared across tree topologies.

We additionally compared SBS spectra produced using the same sequence alignment and maximum likelihood phylogenetic tree but employing either the GTR or HKY models of nucleotide substitution for ancestral reconstruction. These analyses were carried out using previously published alignments and phylogenetic trees of *S. agalactiae* CC1 and MERS coronavirus [18,19].

## Declarations

### Ethics approval and consent to participate

Not applicable

### Consent for publication

Not applicable

### Availability of data and materials

The datasets generated and/or analysed during the current study are available in the GitHub repository at https://github.com/chrisruis/MutTui_manuscript.

### Authors’ contributions

CR, RAF and JP conceived the project. CR and GTH developed the method and software. CR performed the data analysis. RAF and JP provided supervision and aided in the interpretation of results. CR wrote the paper with contributions from all authors. All authors read and approved the final manuscript.

## Acknowledgements

Many thanks to Lauren Bell for designing the MutTui logo.

## References

1. Smith EC. The not-so-infinite malleability of RNA viruses: Viral and cellular determinants of RNA virus mutation rates. PLoS Pathog. 2017;13:e1006254.

2. Duncan BK, Miller JH. Mutagenic deamination of cytosine residues in DNA. Nature. Nature Publishing Group; 1980;287:560–1.

3. Kim K, Calabrese P, Wang S, Qin C, Rao Y, Feng P, et al. The roles of APOBEC-mediated RNA editing in SARS-CoV-2 mutations, replication and fitness. Sci Rep. Nature Publishing Group; 2022;12:14972.

4. Pfaller CK, George CX, Samuel CE. Adenosine Deaminases Acting on RNA (ADARs) and Viral Infections. Annual Review of Virology. 2021;8:239–64.

5. Xia J, Chiu L-Y, Nehring RB, Núñez MAB, Mei Q, Perez M, et al. Bacteria-to-human protein networks reveal origins of endogenous DNA damage. Cell. 2019;176:127-143.e24.

6. Miyahara E, Nishie M, Takumi S, Miyanohara H, Nishi J, Yoshiie K, et al. Environmental mutagens may be implicated in the emergence of drug-resistant microorganisms. FEMS Microbiology Letters. 2011;317:109–16.

7. Wozniak KJ, Simmons LA. Bacterial DNA excision repair pathways. Nat Rev Microbiol. Nature Publishing Group; 2022;1–13.

8. Fukui K. DNA Mismatch Repair in Eukaryotes and Bacteria [Internet]. Journal of Nucleic Acids. Hindawi; 2010 [cited 2020 Nov 18]. p. e260512. Available from: https://www.hindawi.com/journals/jna/2010/260512/

9. Kucab JE, Zou X, Morganella S, Joel M, Nanda AS, Nagy E, et al. A Compendium of Mutational Signatures of Environmental Agents. Cell. 2019;177:821-836.e16.

10. Degasperi A, Zou X, Dias Amarante T, Martinez-Martinez A, Koh GCC, Dias JML, et al. Substitution mutational signatures in whole-genome–sequenced cancers in the UK population. Science. American Association for the Advancement of Science; 376:abl9283.

11. Alexandrov LB, Nik-Zainal S, Wedge DC, Aparicio SAJR, Behjati S, Biankin AV, et al. Signatures of mutational processes in human cancer. Nature. 2013;500:415–21.

12. Nik-Zainal S, Alexandrov LB, Wedge DC, Van Loo P, Greenman CD, Raine K, et al. Mutational Processes Molding the Genomes of 21 Breast Cancers. Cell. 2012;149:979–93.

13. Alexandrov LB, Kim J, Haradhvala NJ, Huang MN, Tian Ng AW, Wu Y, et al. The repertoire of mutational signatures in human cancer. Nature. Nature Publishing Group; 2020;578:94–101.

14. Zou X, Koh GCC, Nanda AS, Degasperi A, Urgo K, Roumeliotis TI, et al. A systematic CRISPR screen defines mutational mechanisms underpinning signatures caused by replication errors and endogenous DNA damage. Nat Cancer. Nature Publishing Group; 2021;2:643–57.

15. Drost J, van Boxtel R, Blokzijl F, Mizutani T, Sasaki N, Sasselli V, et al. Use of CRISPR-modified human stem cell organoids to study the origin of mutational signatures in cancer. Science. American Association for the Advancement of Science; 2017;358:234–8.

16. Alexandrov LB, Ju YS, Haase K, Loo PV, Martincorena I, Nik-Zainal S, et al. Mutational signatures associated with tobacco smoking in human cancer. Science. 2016;354:618–22.

17. Ruis C, Bryant JM, Bell SC, Thomson R, Davidson RM, Hasan NA, et al. Dissemination of Mycobacterium abscessus via global transmission networks. Nat Microbiol. 2021;1–10.

18. Ruis C, Weimann A, Tonkin-Hill G, Pandurangan AP, Matuszewska M, Murray GGR, et al. Mutational spectra analysis reveals bacterial niche and transmission routes [Internet]. bioRxiv; 2022 [cited 2022 Sep 6]. p. 2022.07.13.499881. Available from: https://www.biorxiv.org/content/10.1101/2022.07.13.499881v1

19. Ruis C, Peacock TP, Polo LM, Masone D, Alvarez MS, Hinrichs AS, et al. Mutational spectra distinguish SARS-CoV-2 replication niches [Internet]. bioRxiv; 2022 [cited 2022 Sep 29]. p. 2022.09.27.509649. Available from: https://www.biorxiv.org/content/10.1101/2022.09.27.509649v1

20. Sanderson T, Hisner R, Donovan-Banfield I, Peacock T, Ruis C. Identification of a molnupiravir-associated mutational signature in SARS-CoV-2 sequencing databases [Internet]. medRxiv; 2023 [cited 2023 Mar 1]. p. 2023.01.26.23284998. Available from: https://www.medrxiv.org/content/10.1101/2023.01.26.23284998v1

21. Tonkin-Hill G, Martincorena I, Amato R, Lawson AR, Gerstung M, Johnston I, et al. Patterns of within-host genetic diversity in SARS-CoV-2. Neher RA, Sawyer SL, Neher RA, Lauring AS, editors. eLife. eLife Sciences Publications, Ltd; 2021;10:e66857.

22. Bloom JD, Beichman AC, Neher RA, Harris K. Evolution of the SARS-CoV-2 mutational spectrum [Internet]. bioRxiv; 2022 [cited 2022 Dec 5]. p. 2022.11.19.517207. Available from: https://www.biorxiv.org/content/10.1101/2022.11.19.517207v1

23. Tonkin-Hill G, Ling C, Chaguza C, Salter SJ, Hinfonthong P, Nikolaou E, et al. Pneumococcal within-host diversity during colonization, transmission and treatment. Nat Microbiol. Nature Publishing Group; 2022;7:1791–804.

24. Murray GGR, Balmer AJ, Herbert J, Hadjirin NF, Kemp CL, Matuszewska M, et al. Mutation rate dynamics reflect ecological change in an emerging zoonotic pathogen. PLOS Genetics. Public Library of Science; 2021;17:e1009864.

25. Sagulenko P, Puller V, Neher RA. TreeTime: Maximum-likelihood phylodynamic analysis. Virus Evolution. 2018;4:vex042.

26. Bharati BK, Gowder M, Zheng F, Alzoubi K, Svetlov V, Kamarthapu V, et al. Crucial role and mechanism of transcription-coupled DNA repair in bacteria. Nature. Nature Publishing Group; 2022;604:152–9.

27. Rambaut A, Lam TT, Max Carvalho L, Pybus OG. Exploring the temporal structure of heterochronous sequences using TempEst (formerly Path-O-Gen). Virus Evol [Internet]. Oxford Academic; 2016 [cited 2020 Feb 27];2. Available from: https://academic.oup.com/ve/article/2/1/vew007/1753488

28. Yang Y, Park S-H, Alford-Zappala M, Lee H-W, Li J, Cunningham RP, et al. Role of Endonuclease III Enzymes in Uracil Repair. Mutat Res. 2019;813:20–30.

29. Benjak A, Avanzi C, Singh P, Loiseau C, Girma S, Busso P, et al. Phylogenomics and antimicrobial resistance of the leprosy bacillus Mycobacterium leprae. Nature Communications. Nature Publishing Group; 2018;9:352.

30. Bryant JM, Brown KP, Burbaud S, Everall I, Belardinelli JM, Rodriguez-Rincon D, et al. Stepwise pathogenic evolution of Mycobacterium abscessus. Science [Internet]. American Association for the Advancement of Science; 2021 [cited 2021 Jun 3];372. Available from: https://science.sciencemag.org/content/372/6541/eabb8699

31. Luo T, Xu P, Zhang Y, Porter JL, Ghanem M, Liu Q, et al. Population genomics provides insights into the evolution and adaptation to humans of the waterborne pathogen Mycobacterium kansasii. Nat Commun. 2021;12:2491.

32. Bryant JM, Grogono DM, Rodriguez-Rincon D, Everall I, Brown KP, Moreno P, et al. Emergence and spread of a human-transmissible multidrug-resistant nontuberculous mycobacterium. Science. 2016;354:751–7.

33. Tonkin-Hill G, Lees JA, Bentley SD, Frost SDW, Corander J. Fast hierarchical Bayesian analysis of population structure. Nucleic Acids Res. Oxford Academic; 2019;47:5539–49.

34. Rambaut. A. FigTree. http://tree.bio.ed.ac.uk/software/figtree/ [Internet]. Available from: http://tree.bio.ed.ac.uk/software/figtree/

35. Ronquist F, Teslenko M, van der Mark P, Ayres DL, Darling A, Höhna S, et al. MrBayes 3.2: Efficient Bayesian Phylogenetic Inference and Model Choice Across a Large Model Space. Systematic Biology. 2012;61:539–42.

36. Rambaut A, Drummond AJ, Xie D, Baele G, Suchard MA. Posterior Summarization in Bayesian Phylogenetics Using Tracer 1.7. Syst Biol. Oxford Academic; 2018;67:901–4.

