## Supplementary figures for "Calculating and applying pathogen mutational spectra using MutTui"

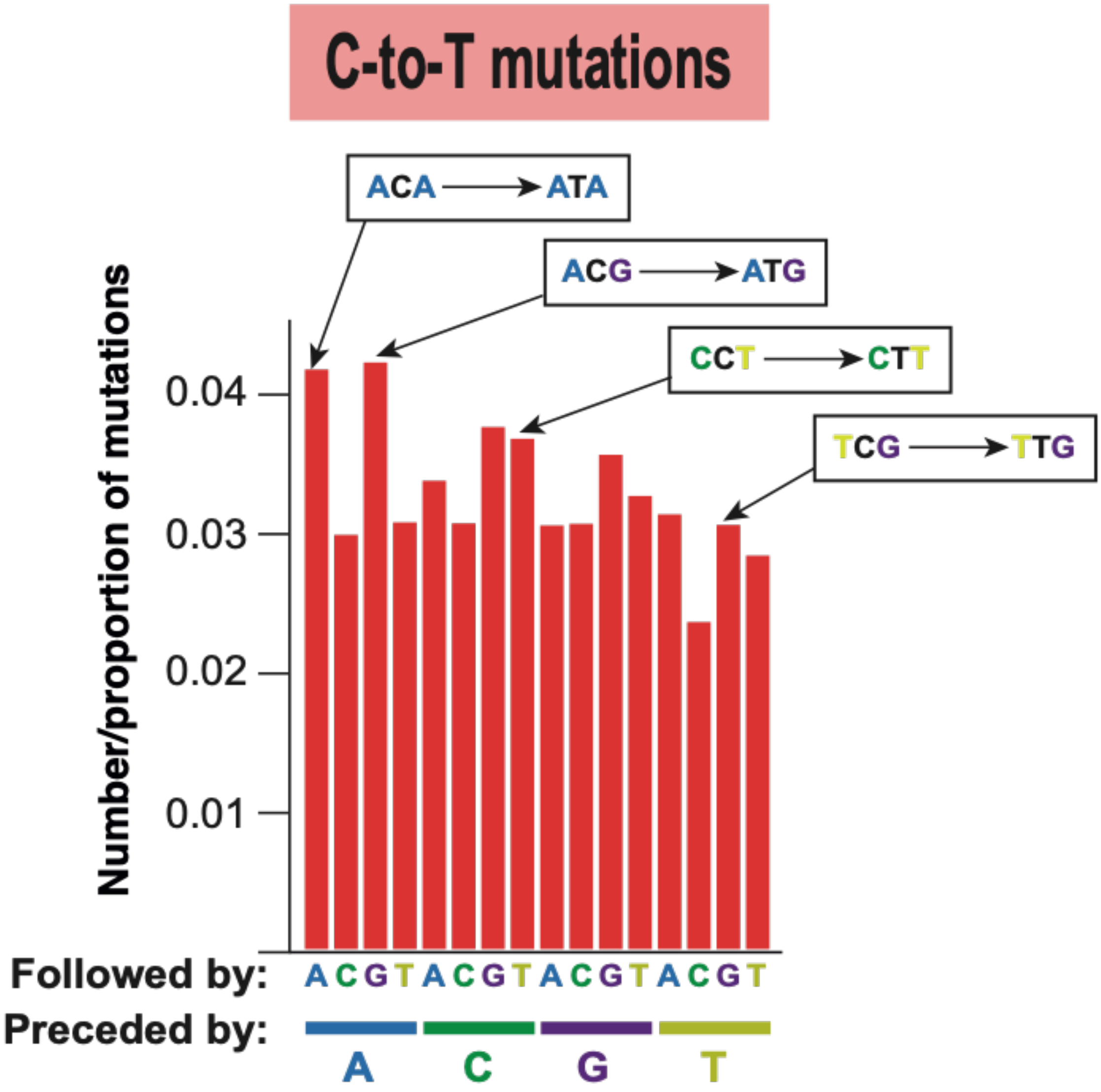


**Figure S1. Summary of contextual mutations in SBS mutational spectra.** In the SBS mutational spectrum, each mutation type is split into 16 surrounding nucleotide contexts, each consisting of the immediately upstream nucleotide and the immediately downstream nucleotide. C-to-T mutations are shown here as an example with the position of each context shown below. Boxed mutations show example contextual mutations.

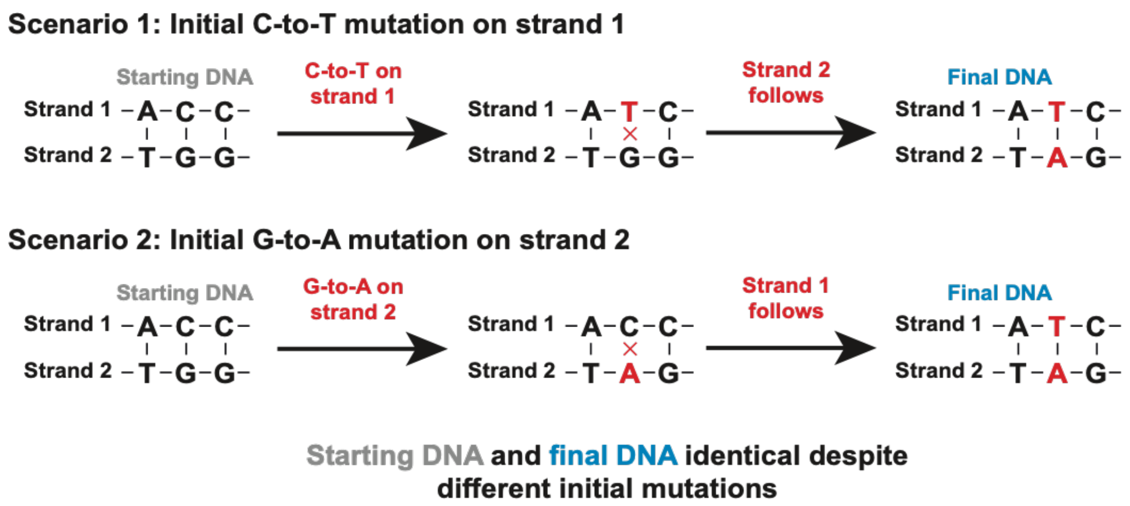


**Figure S2. Symmetric mutations are combined in DNA and dsRNA pathogens.** Different initial mutations on opposing nucleic acid strands can result in indistinguishable mutations. An example is shown depicting 2 scenarios in which the starting and final DNA sequences are indistinguishable despite different initial mutations. MutTui therefore combines symmetrical mutations when calculating mutational spectra for DNA and dsRNA pathogens, for example A[C>T]C and G[G>A]T mutations shown in this example are combined.

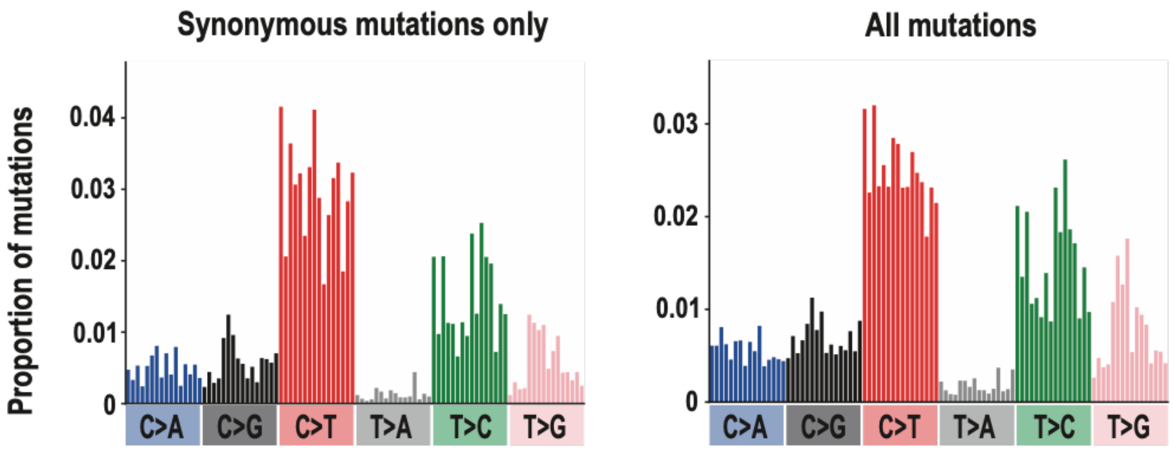


**Figure S3. MutTui can calculate mutational spectra containing synonymous mutations only.** *M. tuberculosis* lineage 4 is shown as an example comparing the SBS spectrum containing synonymous mutations only (providing the --synonymous option and a GFF file to identify genes) with the SBS spectrum calculated using all mutations. Both spectra are rescaled by genomic composition. The SBS spectra are highly similar (cosine similarity 0.98).

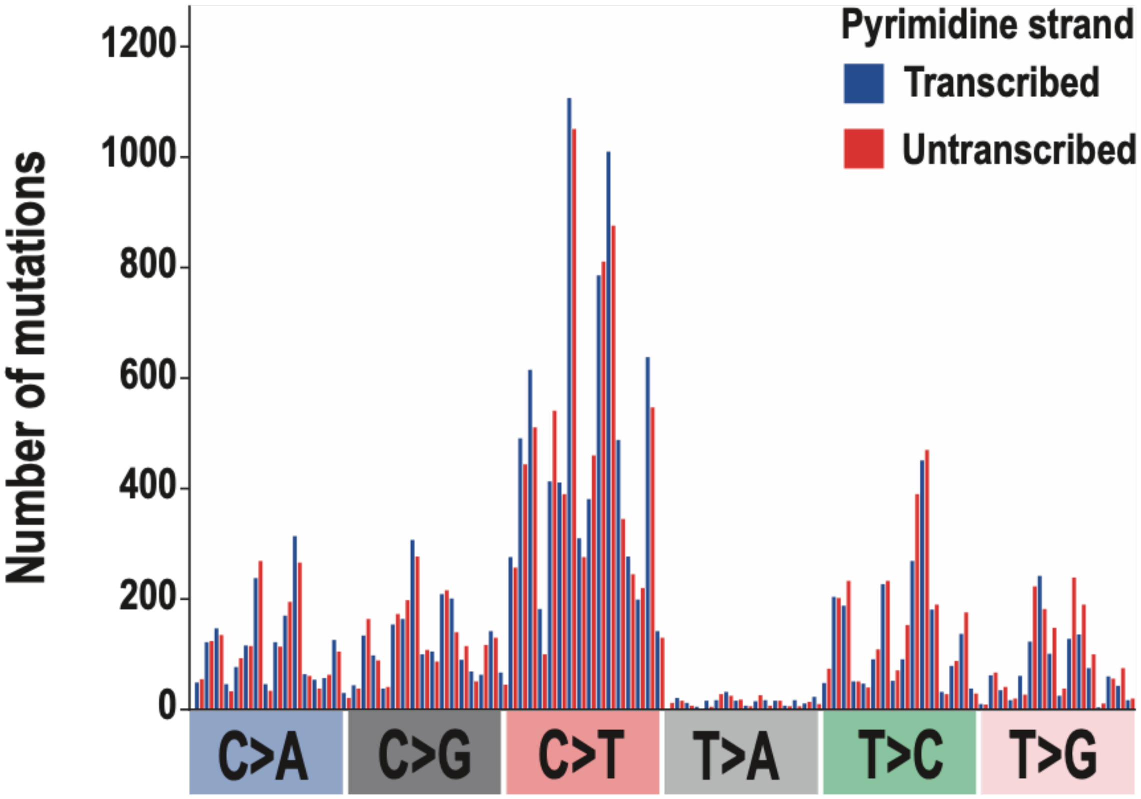


**Figure S4. MutTui can calculate strand bias spectra for DNA and dsRNA pathogens.** *M. tuberculosis* lineage 4 is shown as an example. The strand bias spectrum (calculated by using option --strand_bias and providing a GFF file to identify genes) splits coding mutations based on the strand of the pyrimidine nucleotide. Non-coding mutations are excluded. Within each mutation type, each of the 16 surrounding nucleotide contexts has two adjacent bars showing the count of mutations with the pyrimidine on the transcribed strand and the count of mutations where the pyrimidine is on the untranscribed strand.

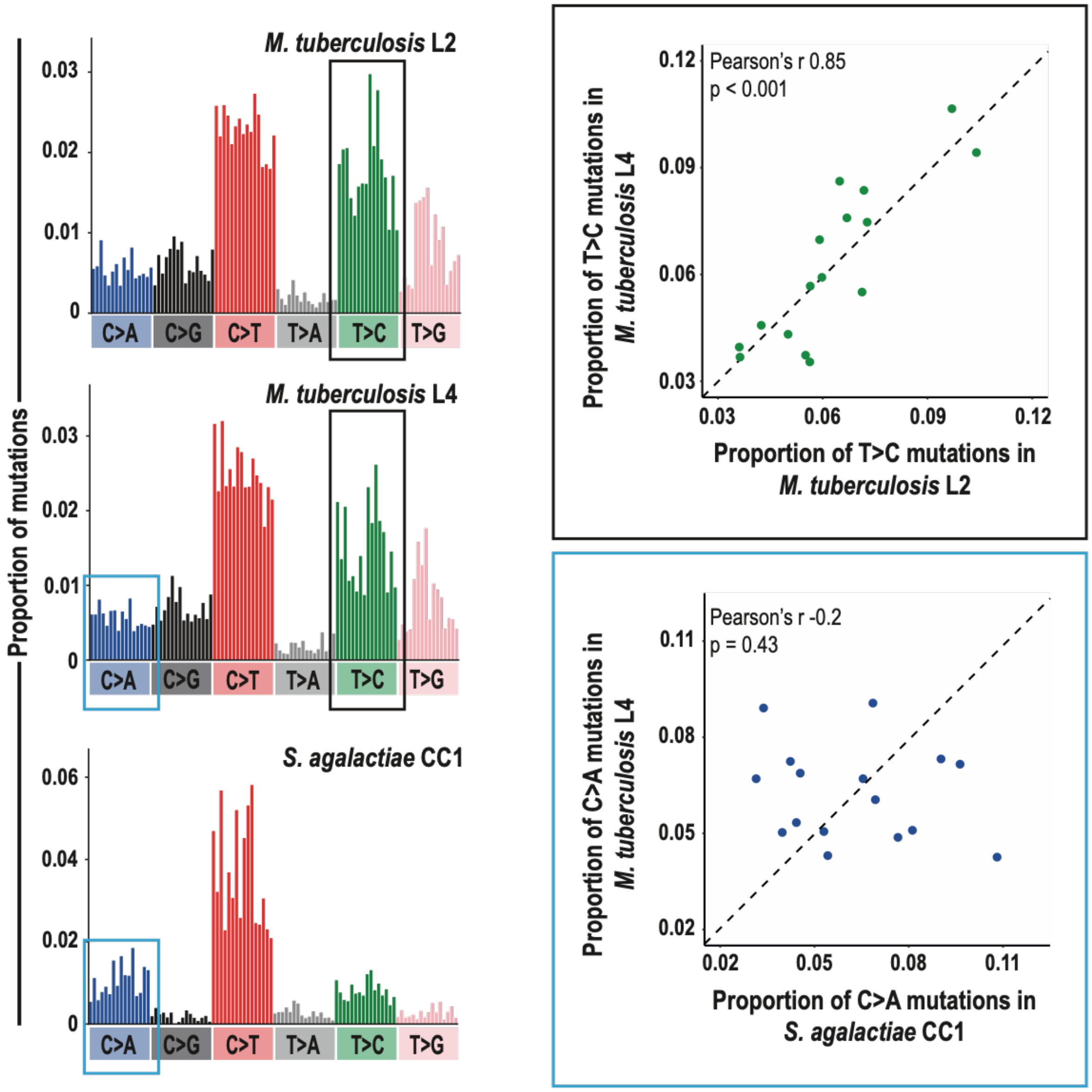


**Figure S5. Comparison of contextual patterns within a mutation type.** MutTui calculates the Pearson's r correlation coefficient comparing contextual mutation proportions. To assess significance, this is compared with the distribution of Pearson's r values across 1000 randomisation of the proportions within each spectrum. Two examples are shown: a significant correlation between the contextual patterns within T>C mutations in the *M. tuberculosis* lineage 2 and lineage 4 spectra (black box); a non-significant correlation between the contextual patterns within C>A mutations in the *M. tuberculosis* lineage 4 and *S. agalactiae* CC1 spectra. The respective full SBS spectra are shown in the left panel with the compared mutation types indicated by rectangles. The dashed lines in the right panels show x=y.

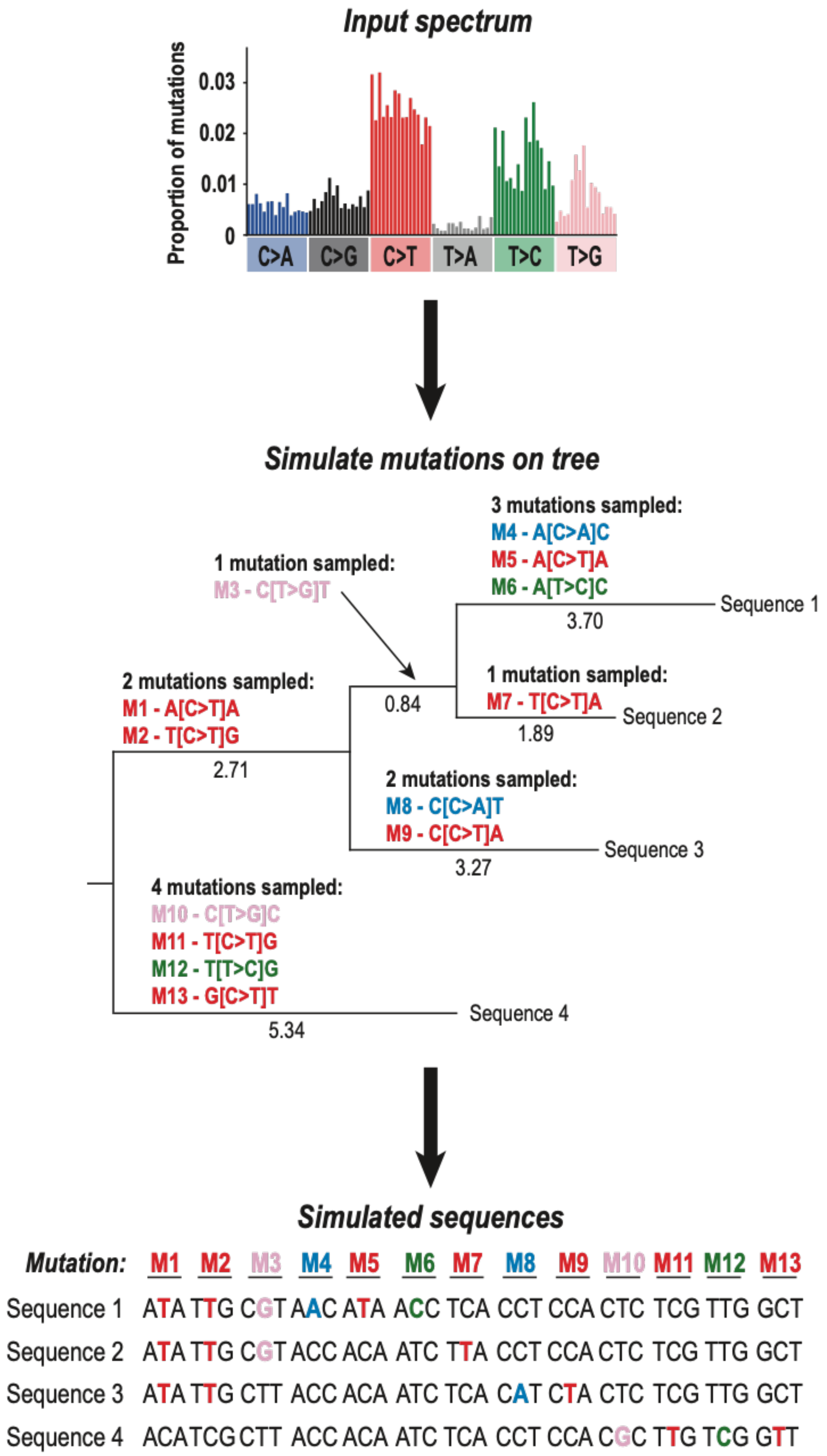


**Figure S6. Overview of simulation procedure to test MutTui.** We simulate mutations under an input SBS spectrum, in this case the SBS spectrum of *M. tuberculosis* lineage 4. Mutations are simulated onto a given phylogenetic tree whose branch lengths are rescaled to sum to the number of target mutations (10,000 for the simulations here). We iterate through the tree and sample mutations on each branch; the number of mutations is a random number from a Poisson distribution with lambda equal to the branch length. This number of mutations is sampled randomly from the input spectrum. Simulated sequences are constructed from each of the simulated mutations; sequences either contain the ancestral triplet if they are not downstream of the mutation, or the mutated triplet if they are downstream of the mutation. MutTui is run on the resulting simulated sequences and the original phylogenetic tree and the calculated spectrum compared to the mutation counts from the simulation.

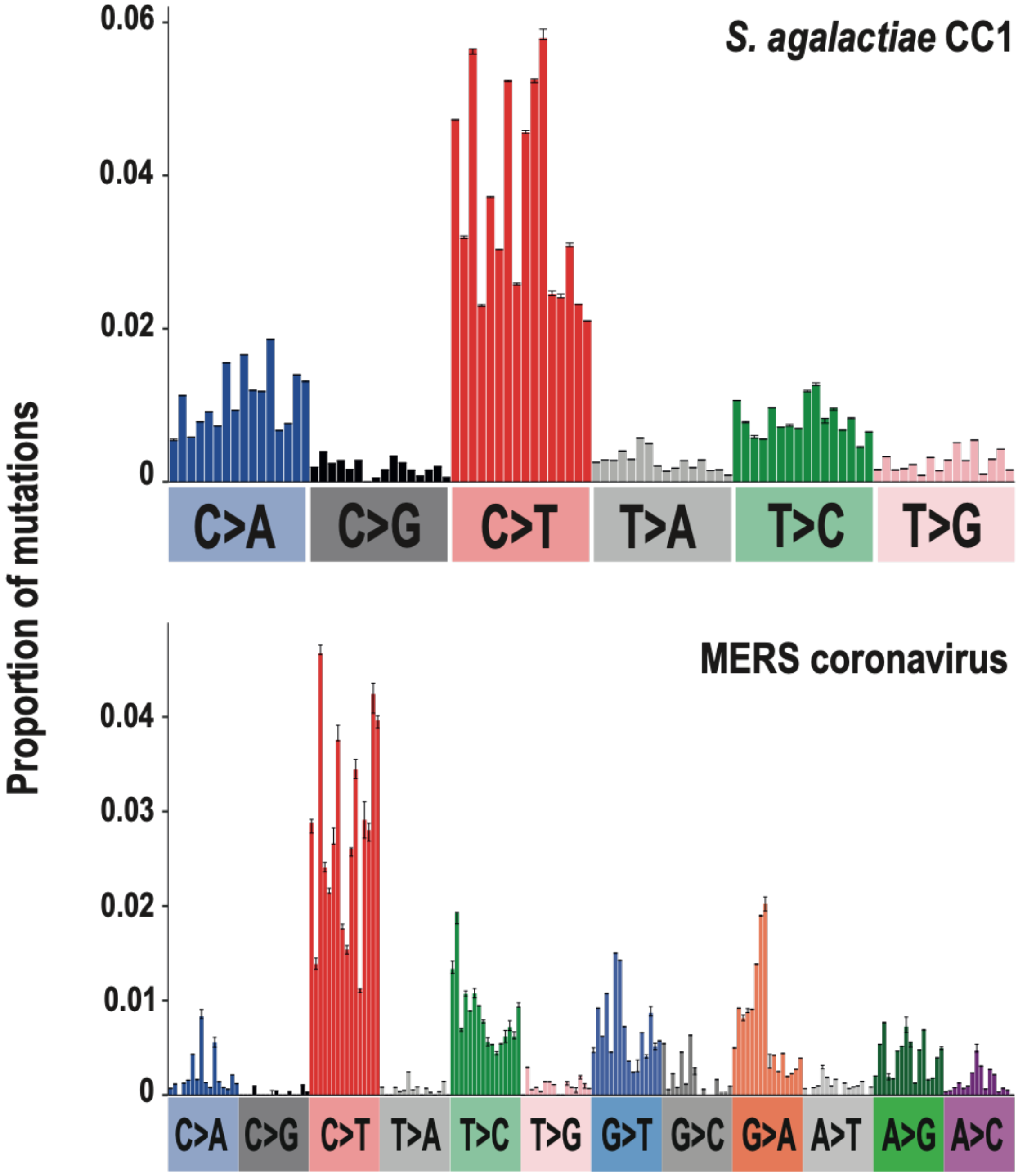


**Figure S7. SBS mutational spectra are robust to tree topology.** We ran MutTui on *S. agalactiae* CC1 and MERS coronavirus datasets with ten alternative tree topologies sampled from a posterior distribution. Bars represent the median mutation proportion across the ten runs and error bars represent the minimum and maximum mutation proportions. Mutation proportions are highly similar across runs.

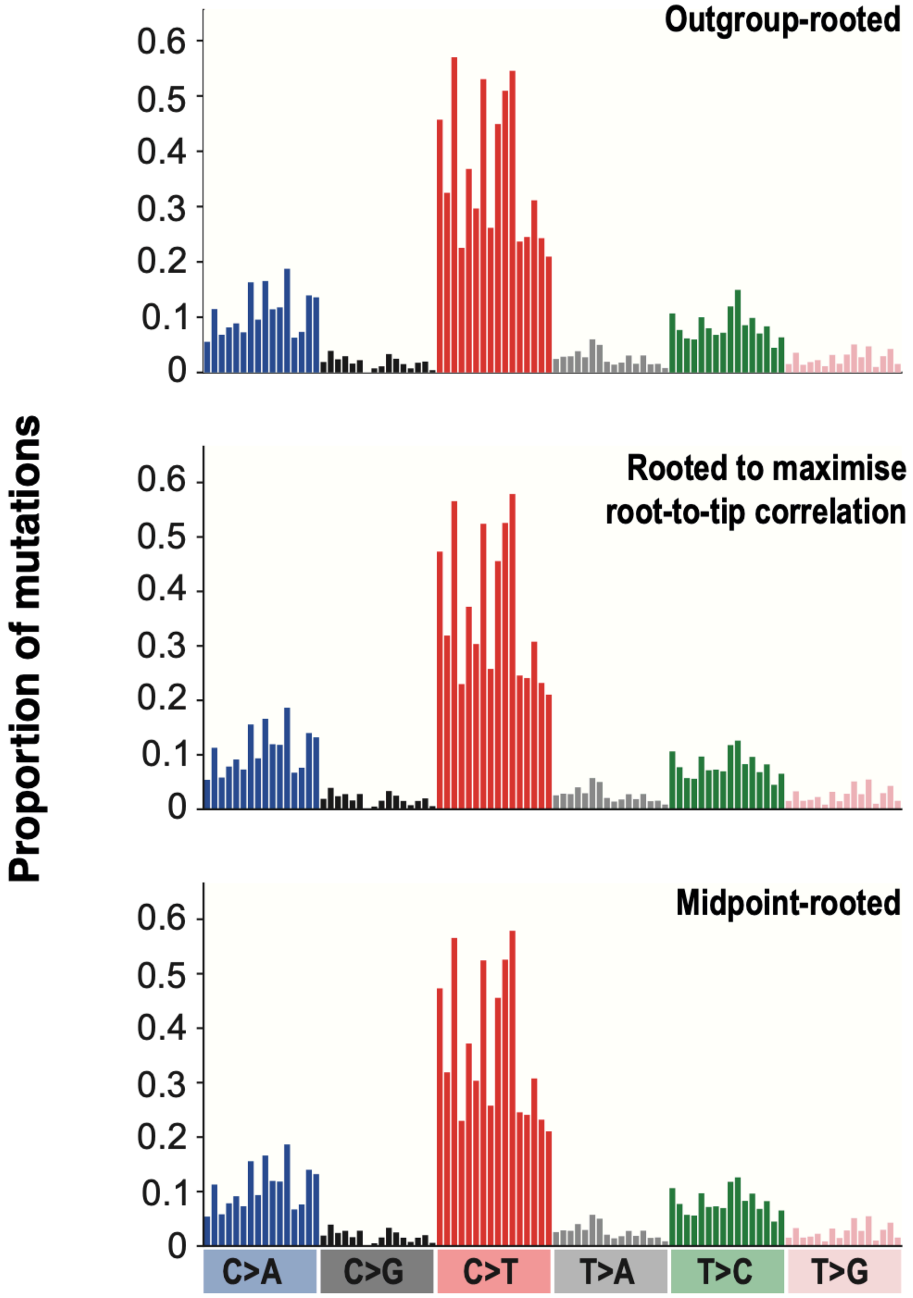


**Figure S8. Comparison of SBS spectra calculated from phylogenetic trees with different rooting strategies.** The *S. agalactiae* CC1 phylogenetic tree was rooted either with a closely related outgroup (top panel), to maximise the heuristic residual mean squared function comparing root-to-tip distance with sample collection date (middle panel) or at the midpoint (bottom panel). MutTui was run using each rooted tree and the same sequence alignment. The resulting SBS spectra are highly similar (cosine similarity >0.99 in each case).

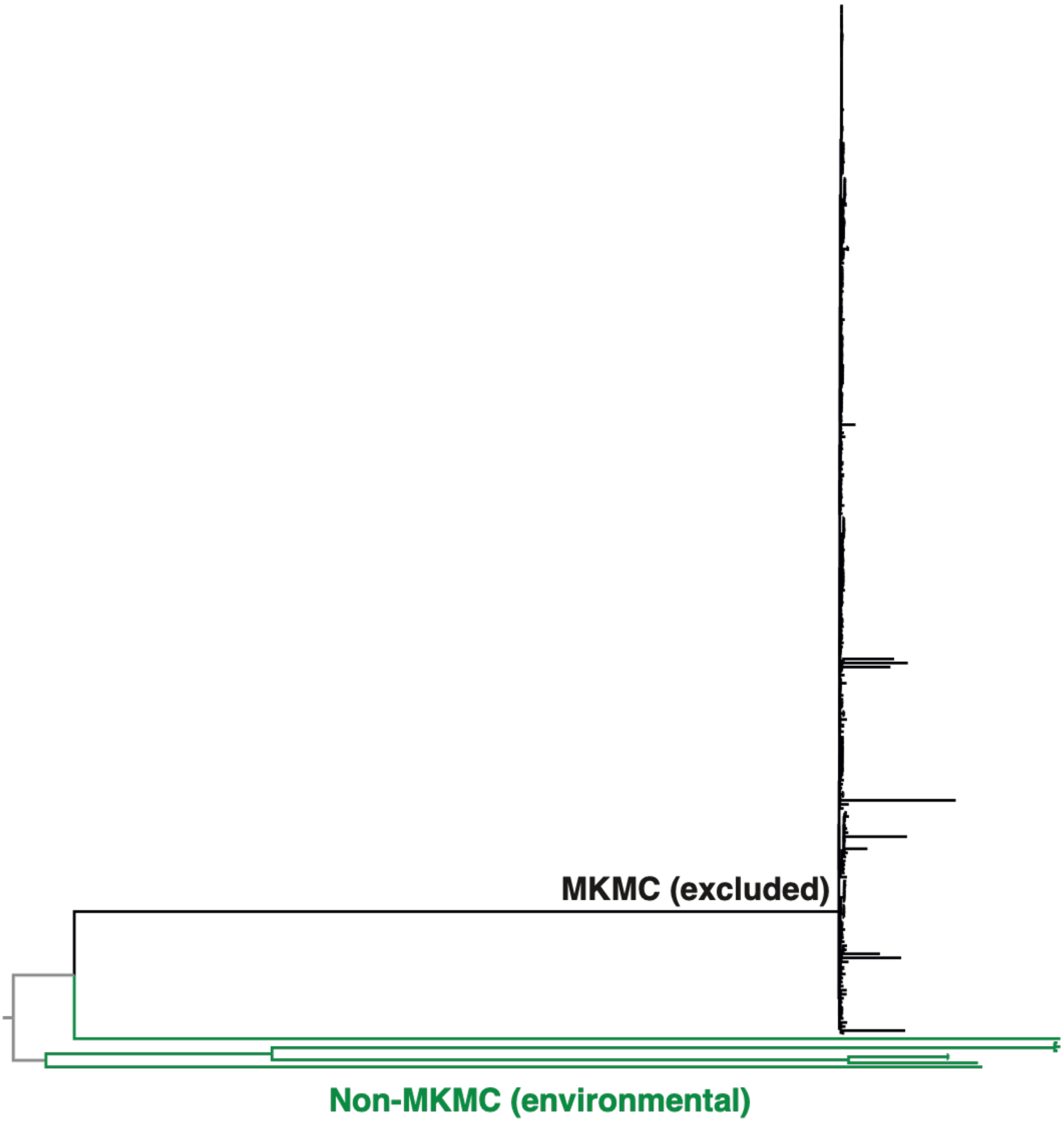


**Figure S9. Classification of phylogenetic branches for calculation of the environmental *M. kansasii* SBS spectrum.** The tree can be divided into the MKMC and non-MKMC. As the niche of the MKMC is poorly understood, we excluded these branches from spectrum calculation. The environmental *M. kansasii* spectrum was calculated from mutations on the non-MKMC branches.

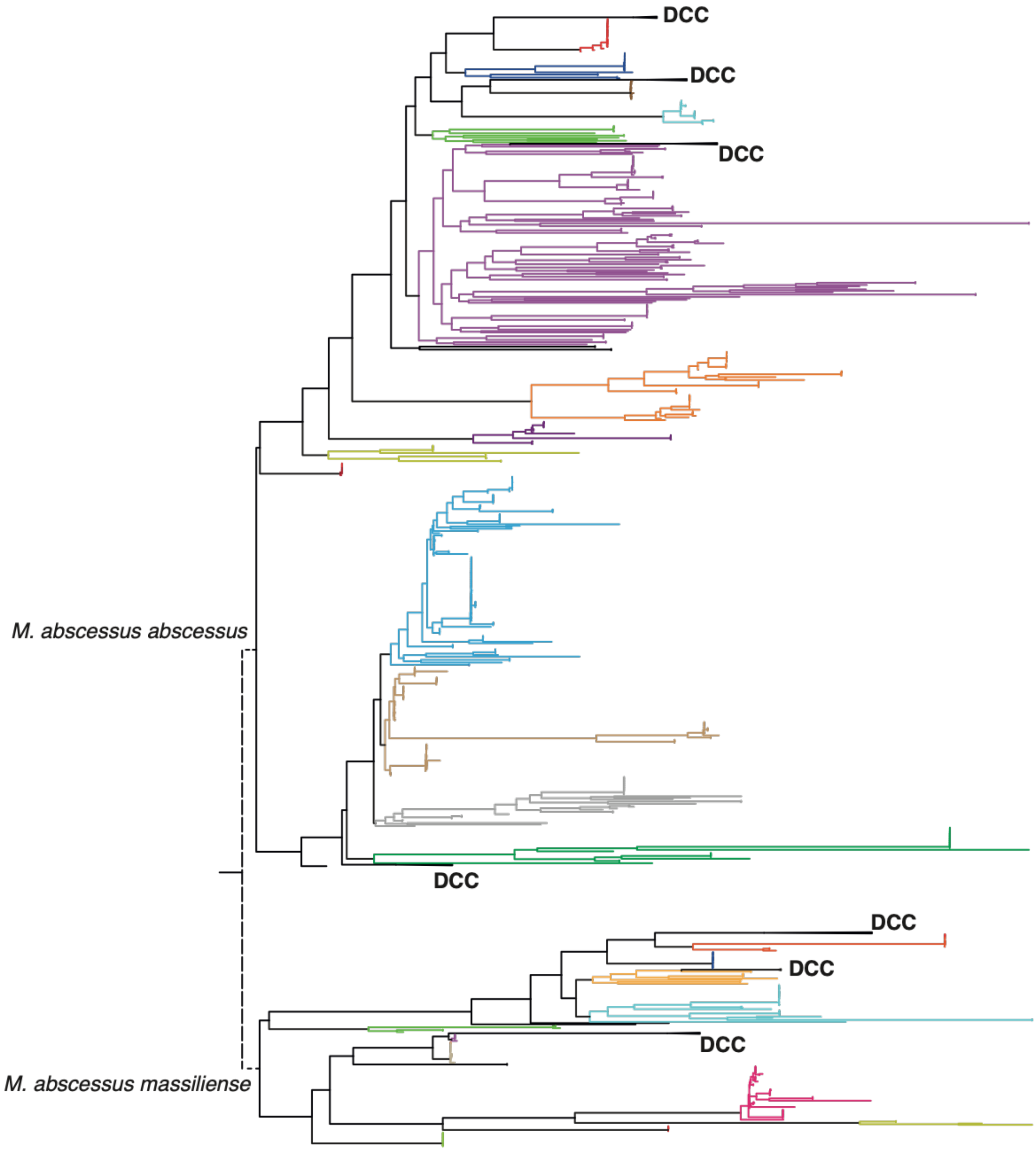


**Figure S10. Clades used in calculation of the environmental *M. abscessus* SBS spectrum.** Due to the high level of diversity within the *M. abscessus* species, we split the dataset into FastBAPS clusters. The dominant circulating clones (DCCs) likely replicate predominantly within the human lung so we did not include these clades in calculation of the environmental *M. abscessus* spectrum; the DCCs are collapsed and shown in black. An SBS spectrum was calculated for each of the remaining clusters separately; each cluster is shown in a different colour. The SBS spectra were subsequently combined to form a single environmental *M. abscessus* spectrum. The two *M. abscessus* subspecies are labelled at the root of the respective clades.
